# Zika: An ongoing threat to women and infants

**DOI:** 10.1101/220640

**Authors:** Beatriz Macedo Coimbra, Flávio Codeço Coelho, Margaret Armstrong, Valeria Saraceni, Cristina Lemos

**Affiliations:** Applied Mathematics School of Getulio Vargas Foundation, Rio de Janeiro, Brazil; MINES Paristech, PSL Research University, Paris, France; Secretaria Municipal de Saúde, Rio de Janeiro, Brazil

## Abstract

Recent data from Rio de Janeiro shows a sharp drop in the number of notified cases of Zika in the summer of 2016-17, compared to the previous summer. This is probably due to herd immunity built up after the previous year's epidemic. There is still a much higher incidence among women than men, almost certainly due to sexual transmission. An unexpected feature of the new data is that there are proportionally far more cases in children under 15 months than in older age classes. By comparing the incidence for 2016-17 with that of 2015-16, we can deduce the proportion of reported cases for men and women, and also verify that the disparity of incidence between them is still present. Women and children still represent risk groups with regard to *Zika* infection, even during a non-epidemic season.

## Introduction

Although Zika is officially no longer an emergency in many Latin-American countries, where it was epidemic in the summer of 2015-16, the threat is not over. In Brazil, a summer came and went without any significant outbreaks. As Cohen (2017) pointed out, the lack of cases this year is probably due to herd immunity. However there were still enough cases (897 cases)in Rio de Janeiro city to compare the incidence in each age class with those for the previous year (Figure 2). The striking features were that (1) there are about 3 times as many cases for women in the sexually active age bracket as for men and (2) there is a peak of cases for infants up to 15 months old who are only now are catching Zika, possibly as a result of the higher risk of infection for women.

Reports from Brazil and Colombia (Pacheco et al., 2016; Coelho et al., 2016) showed up to three times more cases in women than in men in the 2015-16 epidemic. At the time they were published, it was suggested that the excess of female cases could have been due to the state of alert about *Zika* and microcephaly and the increased attention given to cases by health professionals (Maxian et al., 2017). These doubts make it all the more important to carry out new analyses which could either confirm or reject the hypothesis of intrinsic higher risk to women supported by many studies (Hamer et al., 2017; Possas et al., 2017; D’Ortenzio et al., 2016). As there was no epidemic in the summer of 2016-17, people were rightfully relieved. But this also meant we had the perfect opportunity to verify whether there is gender bias in *Zika* incidence, in the absence of a mass scare on the part of women, and of any specific health programs monitoring women for *Zika*.

We start by dissecting the incidence rates by age, gender and mode of transmission in both years, and then proceed to test whether the incidences in women and infants in the non-epidemic period were higher than expected.

## Methods

The dataset used for these analyses was obtained from SINAN, the Brazilian national registry of diseases requiring mandatory reporting. We analyzed all reported cases of *Zika* and dengue, within the city of Rio de Janeiro between September of 2015 and July of 2017. Dengue incidence is used as a control as it shares the same vectorial transmission mechanism with zika but is not sexually transmissible.

For the analysis, we separated the epidemic period from the post-epidemic one. We considered the periods from January 2015 up to August 2016 and September 2016 up to July 2017 as years 1 and 2, respectively.

Let *α*_1_ and *α*_2_ be the proportions of susceptible people who are infected with *ZIKV* via mosquito bite in years 1 and 2, respectively. We assume that these proportions are the same for men and women.

Let *β*_*M*_ and *β*_*W*_ be the proportion of reported *Zika* cases for men and women, respectively. Therefore, the proportion of unreported cases are (1 - *β*_*M*_) and (1 - *β*_*W*_). We also assume that the reporting rates for men and women did not change from one year to the next.

All these rates are outlined in Tables 1 and 2 below.

**Table 1:** Proportions of infected by each route divided by sex and age group in 2016.

|  | Vector transmission |  | Sexual transmission |  | Still susceptible |
| --- | --- | --- | --- | --- | --- |
|  | Reported | Unreported | Reported | Unreported |  |
| Men | $\alpha_1\beta_M$ | $\alpha_1(1 - \beta_M)$ | - | - | $(1 - \alpha_1)$ |
| Women | $\alpha_1\beta_W$ | $\alpha_1(1 - \beta_W)$ | $\gamma_1\beta_W(1 - \alpha_1)$ | $\gamma_1(1 - \beta_W)(1 - \alpha_1)$ | $(1 - \alpha_1)(1 - \gamma_1)$ |

**Table 2:** Proportions of Infected by each route divided by sex and age group in 2017.

|  | Vector transmission |  | Sexual transmission |  | Still susceptible |
| --- | --- | --- | --- | --- | --- |
|  | Reported | Unreported | Reported | Unreported |  |
| Infants | $\alpha_2\beta_W$ | $\alpha_2(1 - \beta_W)$ | - | - | $(1 - \alpha_2)$ |
| Men | $\alpha_2\beta_M$ | $\alpha_2(1 - \beta_M)$ | - | - | $(1 - \alpha_2)$ |
| Women | $\alpha_2\beta_W$ | $\alpha_2(1 - \beta_W)$ | $\gamma_2\beta_W(1 - \alpha_2)$ | $\gamma_2(1 - \beta_W)(1 - \alpha_2)$ | $(1 - \alpha_2)(1 - \gamma_2)$ |

Some other rates can be expressed in terms of the ones presented above. For example the product *α*_1_*β*_*M*_ is the proportion of reported male *Zika* cases due to vector transmission in year 1. As we know that men can only get *Zika* via mosquito bites, we consider all male cases as originating from vector transmission. Thus, as the reporting rate for males (*β*_*M*_) is known to be approximately 0.1 (Bastos et al. Unpublished results) and assuming that the whole population was susceptible to Zika in year 1, to find *α*_1_ we have that

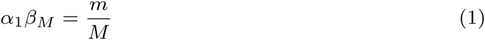

Where *m* is the total reported male cases and *M* is the total male population. Therefore, the rate *α*_1_ can be defined as follows

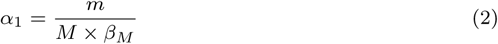

Applying the data into (2), we obtained *α*_1_ ≈ 0.044, meaning that the total number of male Zika cases (including unreported cases) is 133,110.

The proportion of notified female *Zika* cases in year 1, *β*_*W*_ is known to be approximately 0.12 (Bastos et al. Unpublished results).

Let *γ*_1_ and *γ*_2_ be the proportions of susceptible women between 15 and 60 years of age (hereby defined as the sexually active age bracket) who caught *Zika* through sexual transmission in year 1 and 2, respectively. As we consider sexual transmission only in the analysis of female cases, the product *γ*_1_*β*_W_(1 − *α*_1_) is the proportion of notified, sexually contracted, female *Zika* cases in year 1 and, consequently, the calculation with respect to year 2 is analogous. We assume the sexual transmission from women to men to be insignificant (Davidson, 2016).

To assess the excess of cases in women in the sexually active age class, we applied a chi-square test to the case counts for both men and women in the age classes between 15 and 60 years old.

The susceptible population in year 2 is different from the one in year 1. After the first epidemic, we assume that people which had Zika became immune to it, therefore, to determine the susceptible population size for year 2, we deducted Zika cases in year 1 from the total population. As we denote by 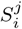 the number of susceptible people from the group *j* in year *i*, we have the following scheme

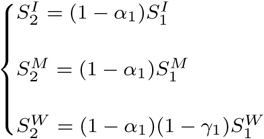

where the groups *I*, *M* and *W* correspond, respectively, to infants, men and women.

Let *α*_2_ be the proportion of susceptible people who catch *Zika* via mosquito bite in year 2. Assuming that this proportion is the same for all age groups, we focus on the group of men, who are not at risk of sexual transmission. Then, the product *α*_2_*β*_*M*_ is the proportion of notified men cases in year 2. Knowing that, we calculated *α*_2_ by dividing the number of cases in men in year 2 by the population of the susceptible ones in the same year multiplied by *β*_*W*_. The number of susceptible men in year 2 was obtained from the same number for year 1, weighted by the rate (1 − *α*_1_), since it represents the proportion of susceptible people for this group. We obtained *α*_2_ ≈ 0.0010.

## Results

Figure 1, shows the incidence time series for both periods. It is clear that there was no outbreak in Rio de Janeiro in year 2, but transmission never stopped completely after the epidemic. For this analysis, we assumed that in the post-epidemic period (2016-17), the risk of getting Zika through the vectorial route is independent of gender and age. This means the rate ck2, which tells us the proportion of susceptible people who get Zika via mosquito bite in year 2, should be the same for children and men (table 2).

**Figure 1:**
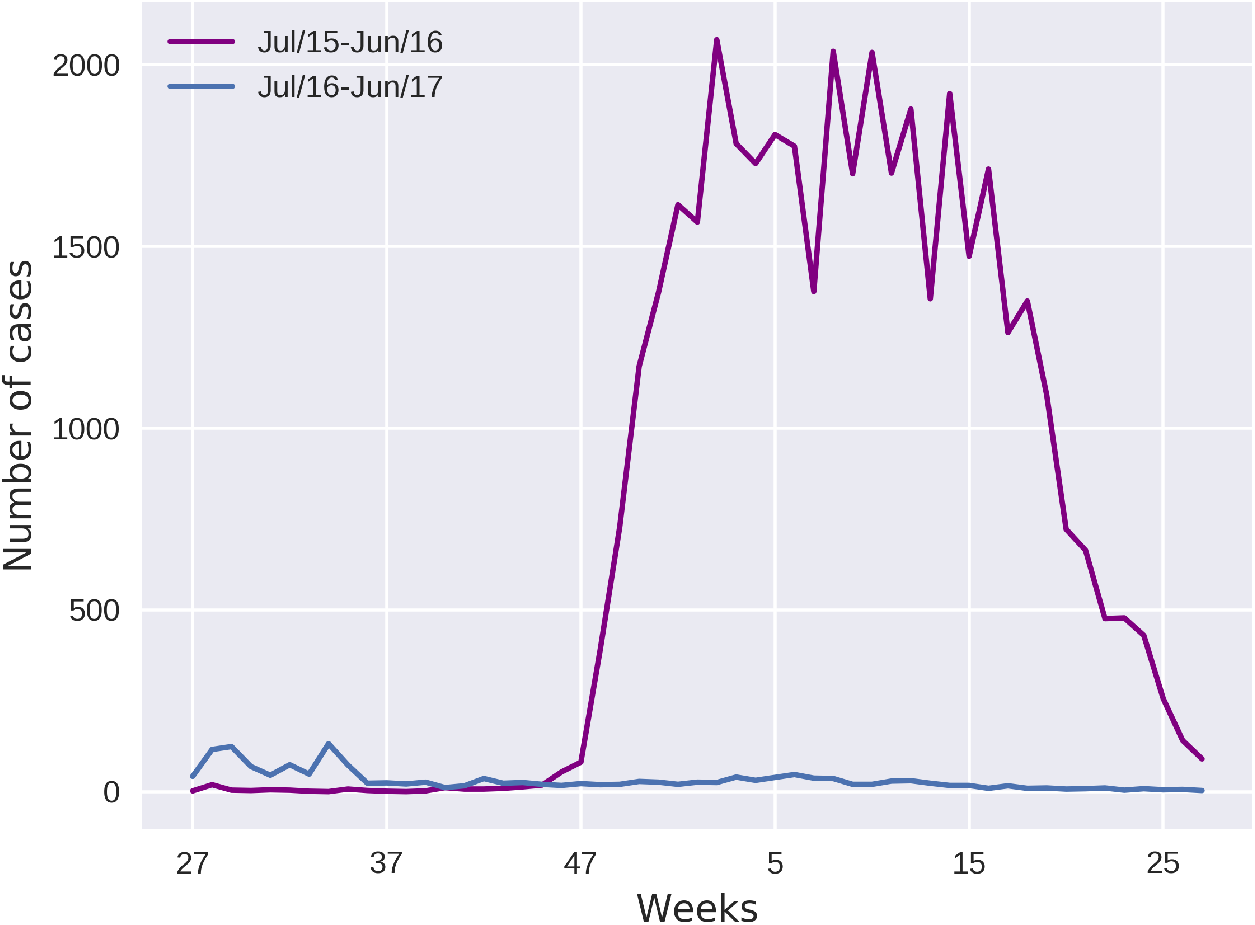
Comparison between the number of cases in both periods. Both Series start on epidemiological week 27, ending on the week 26 of the following year.

Figure 3 shows the age distribution of cases of Zika per week for women and men within Rio de Janeiro city in year 2 (09/2016 to 07/2017). Women between 15 and 60 years old show 3-4 times the incidence of men of the same age (474 women cases against 174 for men). This pattern is similar to that observed in the previous (epidemic) year (figure 2). A chi-squared test confirmed a higher than expected risk of Zika for the female population within the sexually active age bracket (Table 5, *p* < 10^−27^ and *χ*^2^ = 115.77).

**Figure 2:**
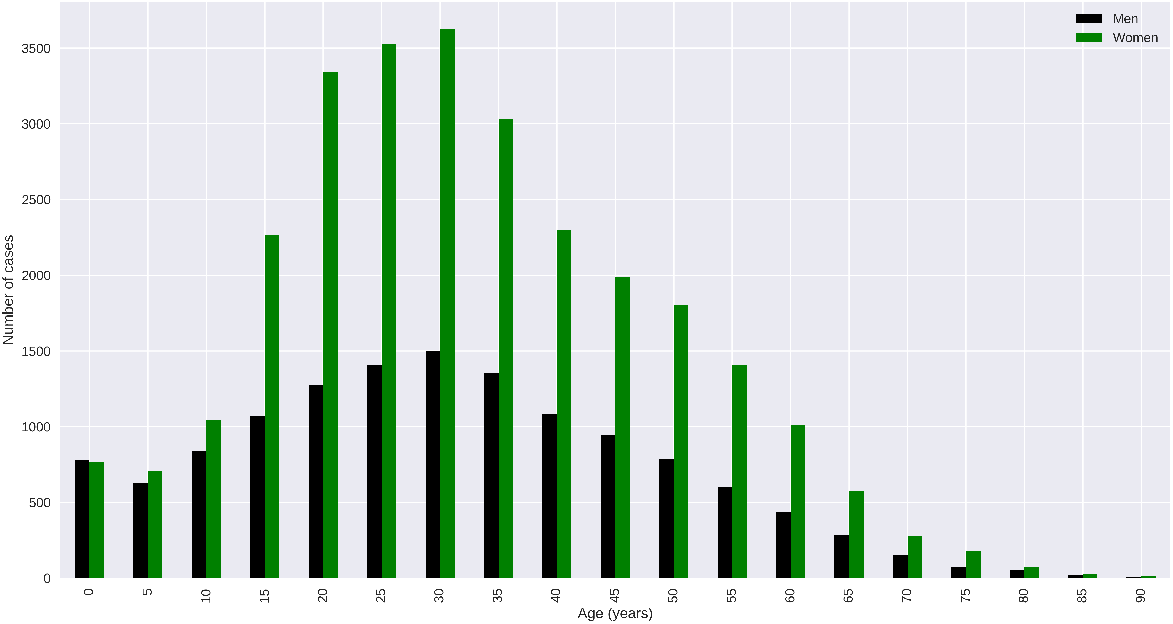
*Zika* cases of Rio de Janeiro from January 2015 to August 2016

**Figure 3:**
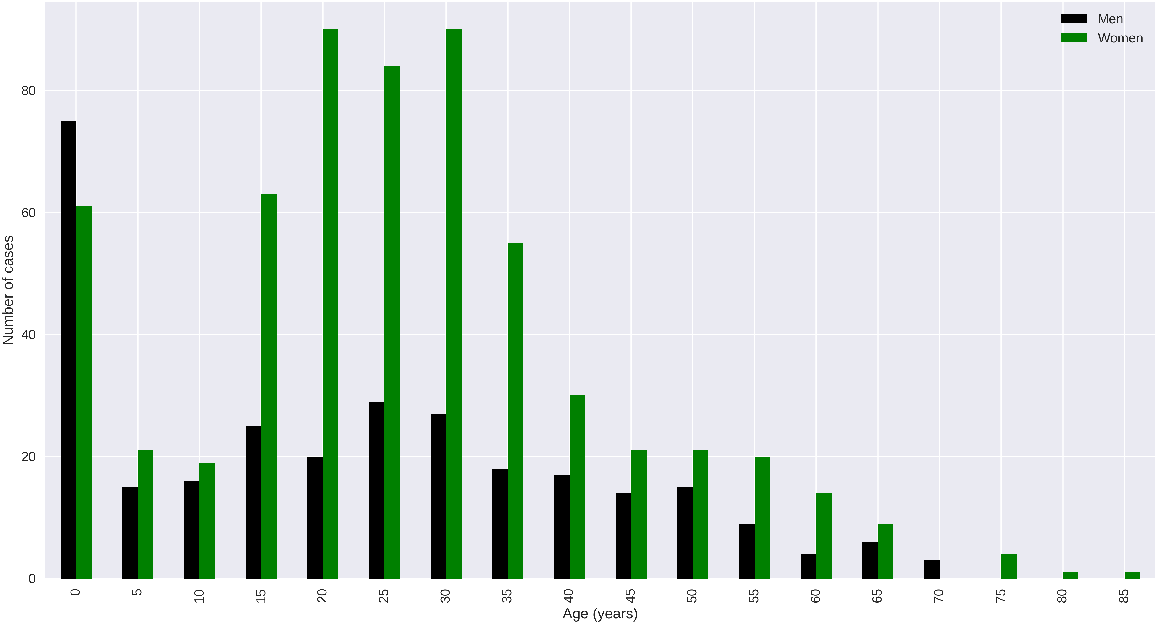
*Zika* cases of Rio de Janeiro from September 2016 to July 2017

If we calculate this rate (*α*_2_) based on the men cases, we obtain *α*_3_ ≈ 0.001. However, if we perform the same calculation based on the cases of children from 0 to 4 years old, we get approximately 0.003. It means that, in fact, children have around 3x more risk of infection of Zika, compared to the group of men.

A chi-squared test confirmed a higher than expected incidence rate of Zika for children younger than 4 years old (Tables 3 and 4). The expected numbers returned by this contingency table are very similar numbers to the ones we obtained from the rates outlined in Tables 2 and 7.

**Table 3:** Observed number of cases in children(between 0 and 4 years old) and adults the population of Rio de Janeiro in 2016-17.

|  | Zika cases | No Zika |
| --- | --- | --- |
| Children(0-4) | 136 | 337,191 |
| Adults | 761 | 6,063,867 |

**Table 4:** Expected number of cases in children (between 0 and 4 years old) and adults for the population of Rio de Janeiro in 2016-17, assuming equal infection rates in both groups.

|  | Zika cases | No Zika |
| --- | --- | --- |
| Children(0-4) | 47 | 337,279 |
| Adults | 849 | 6,063,778 |

**Table 5:** Observed contingency table for the sexually active population (15 to 60 years old) of Rio de Janeiro in 2017.

|  | Zika cases | No Zika |
| --- | --- | --- |
| Men | 174 | 2,011,783 |
| Women | 474 | 2,175,157 |

**Table 6:** Results for Table 1. Numbers for the epidemic year. Percentages of the total susceptible population which got Zika or remained susceptible. In parenthesis are the raw number of cases, based on our estimates for the mentioned rates.

|  | Vector transmission |  | Sexual transmission |  | Still susceptible |
| --- | --- | --- | --- | --- | --- |
|  | Reported | Unreported | Reported | Unreported |  |
| Men | 0.44%(13,311) | 4.00% (119,799) | - | - | 95.55% (2,860,907) |
| Women | 0.53% (18,098) | 3.91% (132,724) | 0.60% (20,356) | 4.40% (149,282) | 90.55% (3,071,963) |

**Table 7:** Results for Table 2. Numbers for the non-epidemic period (September 2016 up to July 2017). Percentages of the total susceptible population which got Zika or reamined susceptible. In parenthesis are the raw number of cases, based on our estimates for the mentioned rates.

|  | Vector transmission |  | Sexual transmission |  | Still susceptible |
| --- | --- | --- | --- | --- | --- |
|  | Reported | Unreported | Reported | Unreported |  |
| Infants | 0.012% (40) | 0.090% (296) | - | - | 99.897% (328,648) |
| Men | 0.010% (275) | 0.092% (2,482) | - | - | 99.897% (2,690,656) |
| Women | 0.012% (358) | 0.090% (2,630) | 0.013% (393) | 0.098% (2,882) | 99.785% (2,912,657) |

Figure 4 shows a histogram of both sexes in this age bracket, and shows that the majority of cases are for children below 15 months of age and thus born after the end of the previous epidemic (99 cases, from a total of 136 of the whole age group).

**Figure 4:**
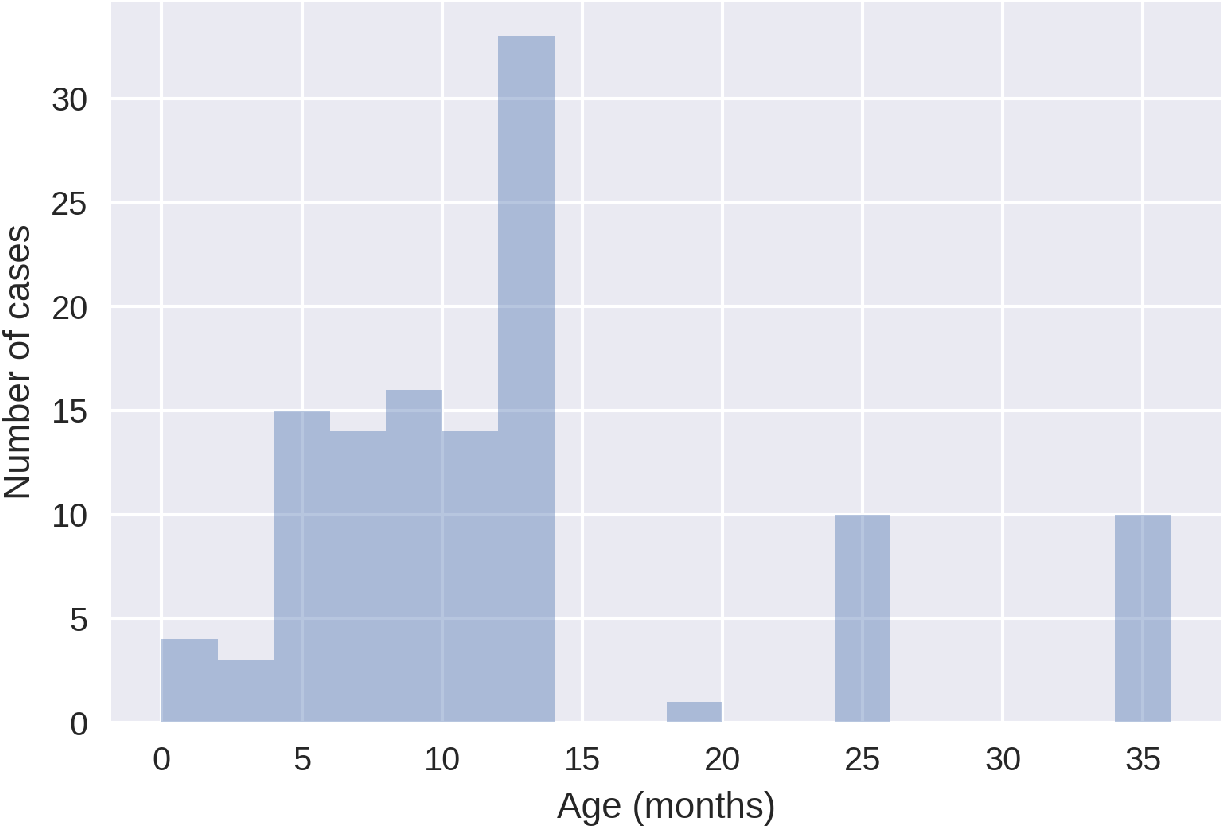
Age distribution of children reported as ZIKV positive in year 2

Finally, to rule out the possibility of the age-related incidence patterns being caused by differential exposure to mosquito bites, we present the age-distributed incidence of dengue for 2016-17 (year 2) on figure 5.

**Figure 5:**
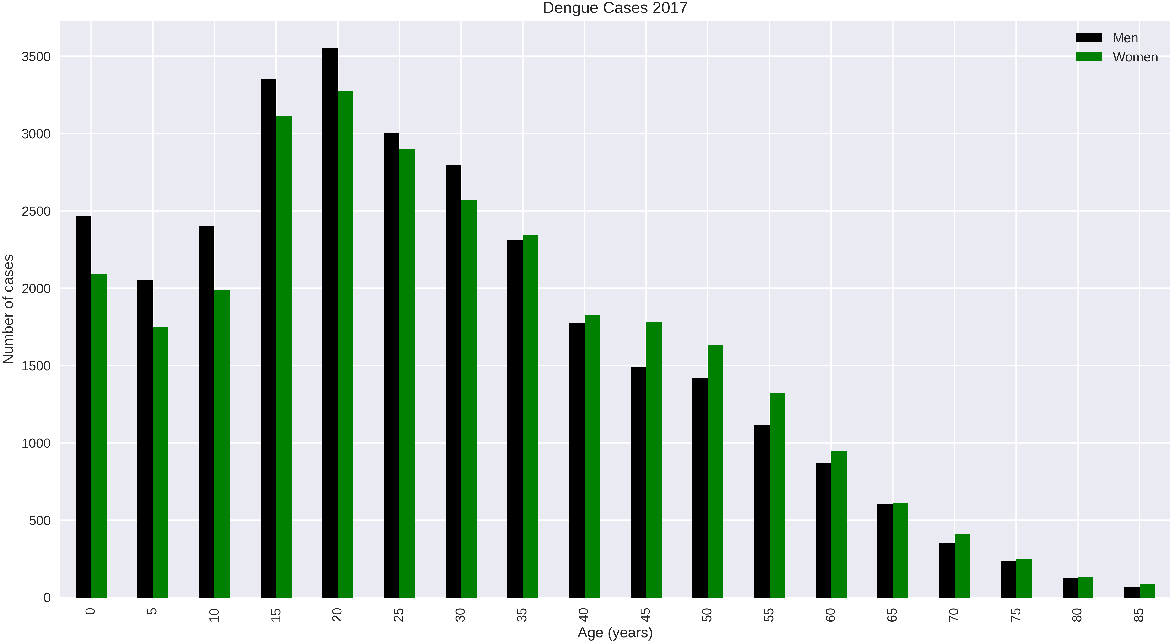
Dengue cases in Rio de Janeiro from September 2016 to July 2017

## Discussion

The difference in incidence between women and men in the age bracket between 15 and 60 years of age confirms our previous results (Coelho et al., 2016) and those from Colombia(Pacheco et al., 2016). It suggests a strong contribution of the sexual transmission route even during non-epidemic periods. The age bracket within which the effect is present is more sharply delimited, possibly due to the reduced intensity of vector transmission in the 2016-17 dataset. A more intense vectorial transmission would tend to attenuate the difference between the incidence in age classes strictly infected by mosquitoes and the sexually active age group.

Here we remark that external effects such as increased female notification due to the emergency status of Zika are not likely to have played a role in the summer of 2016-17. It is also interesting to compare the age-distributed Zika incidence after the last epidemic with that of Dengue for the same period in Rio, which does not show any gender bias. If there were some lingering fear about Zika affecting the diagnostic-seeking behavior of women, we would see some impact in dengue reporting rate for women as well since both disease have very similar clinical manifestations. But if we look at figure 5, no excess female incidence of dengue is seen.

Unlike the age distribution of cases of 2015-16, we observe an excess of cases in children of both sexes under 4 years of age (figure 3). The majority of these cases are under 15 months of age. Due to the low number of cases and the lack of laboratory confirmations, we cannot fully explain this phenomenon, but we hypothesize that children of this age spend most of their time in the presence of women of reproductive ages in the household where the mosquito is ubiquitous, thus the may have been infected by mosquitos which bit their mothers or care-givers at the day-care or pre-schools. The data nevertheless highlights potential risks of Zika for young children in a post epidemic setting.

## Conclusions

Zika remains a serious threat mainly for women and young children. There is still a great deal of uncertainty about the infectious period for sexual transmission in males. This dataset shows, unequivocally that even during non-epidemic periods, the sexual transmission continues to be important and induce a risk of infection almost 4 times higher for women than for men. These extra women cases will continue to influence the risk to infants, with and without serious neurological damage. We must remember that the non-microcephalic Zika babies may still display milder but not necessarily less important issues such as macular atrophy(Ventura et al., 2016), congenital contractures(Moore et al., 2017) and various neurological sequelae.

Our main conclusions are that the protection of women against Zika infection must be redoubled in countries with ongoing transmission, and that more detailed studies should be funded to help answer the open questions about the long term effects of sexual transmission to the potential endemization of Zika and to the consequent increase in the burden caused by this disease.

